# Identifying transcriptomic bias across developmental shifts in insects

**DOI:** 10.64898/2026.06.12.731678

**Authors:** Sarah Cornet, Alice Dennis

## Abstract

**Background:** Synonymous mutations, once considered neutral, can affect translation efficiency through mRNA folding and splicing, generating codon usage bias. This bias is often linked to genomic GC content, which also influences gene regulation. In the parasitoid wasp *Lysiphlebus fabarum*, GC content was previously shown to shift between developmental stages, with larvae showing higher GC than adults. Whether this phenomenon is widespread among insects remains unknown.

**Results:** Transcriptomic data from six insect species spanning Diptera, Hymenoptera, and Lepidoptera was used to compare GC content between expressed genes in larvae and adults. In five species, larval transcripts exhibited higher GC content than adult transcripts. Differential expression analysis revealed that stage-biased genes displayed consistent GC shifts, and orthologous gene families with representatives across species showed particularly GC-rich larval-biased genes in Hymenoptera and Diptera. At the genome scale, modeling in 317 insect species demonstrated an association between parasitic lifestyle and reduced mean GC content in Hymenoptera and Diptera, providing a possible ecological explanation for AT-rich genomes.

**Conclusions:** Our results show that GC content is dynamic across developmental stages, independent of overall genome composition. Stage-specific GC enrichment may reflect adaptive codon usage optimizing translation during energetically demanding life-history stages such as larval development. Furthermore, the association between parasitism and reduced genomic GC highlights how ecological lifestyle might with genome content and evolution. Lastly, this work identifies candidate genes underlying stage-specific GC bias and provides new insights into the interplay between molecular evolution, development, and parasitic adaptation in insects.

## Introduction

Genetic variation provides the substrate for natural selection, enabling organisms to adapt to their environments. Mutations, ranging from single nucleotide substitutions to both large and small indels and inversions, generate diversity, and their evolutionary fate depends on how they affect fitness. A central distinction is between synonymous substitutions, which do not alter the encoded amino acid, and non-synonymous substitutions, which do. The ratio of non-synonymous to synonymous mutations rates (dN/dS) has become a cornerstone of molecular evolution, as it provides a means to infer selective pressures action on protein-coding genes (Kryazhimskiy & Plotkin, 2008).

While often considered neutral, synonymous mutations are not necessarily silent. They can influence gene expression and protein production through mechanisms such as codon usage bias, mRNA secondary structure, splicing, and translation kinetics (Chamary & Hurst, 2005; Drummond & Wilke, 2008; Sarkar et al., 2022). Codon usage bias, the preferential use of specific codons over synonymous alternatives, is shaped by mutation, drift and selection. Optimal codons tend to match abundant tRNAs, thereby enhancing translational efficiency, whereas rare codons may slow elongation or trigger mRNA decay (Hanson & Coller, 2018; Qian et al., 2012). This bias can affect co-translational protein folding, circadian regulation, and gene expression across taxa (Goodman et al., 2013; C.-H. Yu et al., 2015). Codon usage is also linked to GC content bias, a genome-wide feature shaped by mutational patterns and biased gene conversion. High GC levels correlate with gene expression and stability in many eukaryotic organisms, whereas some organisms, including insects, are known to exhibit unusually AT-rich genomes. This raises questions about the adaptive significance of reduced GC content (Assaf et al., 2017; Alice B. Dennis et al., 2020; Kyriacou et al., 2024).

Gene expression regulation is particularly critical during development, where transcriptional programs guide transitions between life stages. In holometabolous insects, metamorphosis involves dramatic morphological and physiological remodeling, orchestrated by hormonal signaling and stage-specific transcription factors (Truman & Riddiford, 2019). Codon bias and GC content may interact with these regulatory processes by modulating expression efficiency of developmentally important genes. Insects also vary widely in the use of regulatory mechanisms such as DNA methylation: while largely absent in Diptera, it remains functional in some Hymenoptera, where it influences expressions of housekeeping and developmental genes (Glastad et al., 2019). How genomic features such as codons bias and GC content influence stage-biased gene expression remains poorly understood, particularly in species with unusually AT-rich genomes.

In addition to developmental constraints, insects face strong selective pressures from ecological interactions. Host-parasitoid systems are a striking example, where coevolutionary dynamics drive adaptive change in both partners (Hembry & Weber, 2020). Parasitoid wasps and flies lay their eggs within or upon a host, and their larval development is fatal to the host (Mackauer et al., 1997). Hosts can mount defenses such as immune encapsulation or protective symbionts (e.g., *Hamiltonella defensa* in aphids), while parasitoids counter-adapt through venoms and immune suppression strategies (Dennis et al., 2017; Wertheim et al., 2005). These antagonistic interactions impose strong selective pressures that can leave genomic signatures, including shifts in gene expression and sequence composition. Understanding how codon bias, GC content, and gene regulation interact in parasitoids may therefore illuminate both developmental biology and coevolutionary adaptation.

Despite extensive work on codon usage, GC bias, and host-parasite interactions, little attention has been paid to these systematically or in relation to developmental stages. We do not know how these forces jointly shape stage-biased gene expression in parasitoid systems. In this study, we investigated larvae- and adult-biased genes, focusing on their codon usage patterns, GC content and enrichment for biological processes. By integrating evolutionary and functional perspectives, we examine how molecular constrains, and ecological pressures could contribute to insect adaptation. We tested three hypotheses: (1) low GC content is associated with parasitism and may provide adaptive advantages; (2) GC shifts across developmental stages improve translational efficiency through codon optimization; and (3) these shifts represent convergent adaptations rather than inheritance from a common ancestor.

## Material and methods

### Data acquisition

The RNA-seq datasets were procured from the NCBI Sequence Read Archive (SRA, accession numbers in Supplementary Material). The species selected for analysis came from the orders Diptera, Hymenoptera and Lepidoptera and were chosen because there were the appropriate life-history stages and sufficient samples to yield high-quality data. These taxa represent well-studied insects, and this taxonomic breadth permitted the examination of species exhibiting diverse methylation mechanisms and life history strategies, which could provide insights into potential adaptations in GC content. Datasets were selected if they differentiated males, females, and larvae, with a minimum of three RNA-seq files per condition and were prioritized to ensure comprehensive representation. Data from *Lysiphlebus fabarum* were obtained from the Supplementary material provided by Dennis *et al*. (2020). The reference genomes of the six species were obtained from the National Center for Biotechnology Information (NCBI) and Ensembl databases (Supplementary Materials).

The whole-genome dataset used for our model contains 317 insect species, obtained from Dennis et al. (2020) and Kyriacou et al. (2024), with 52 additional parasitic species retrieved from the NCBI database to ensure balanced sample sizes for Diptera and Hymenoptera. To prevent duplication, instances where sources overlapped were eliminated.

### RNA-seq analysis

#### Data processing and cleaning

The RNA sequences of the entire organism, with the exception of *Manduca sexta*, which only encompasses the head, were obtained from NCBI using the SRA Toolkit (3.0.5; (SRA Toolkit Development Team, 2023). The quality of the sequences was evaluated using FastQC (0.12.1; Andrews S et al., 2010) both before and after trimming. Trimmomatic (0.39; Bolger et al., 2014) was used to remove short and low-quality reads, the sliding window was set to 4:15 with the minimum length dependent on the size of the species reads, from 20 to 75 bp. This setting evaluates the quality scores over a sliding window of 4 bases and removes sequences where the average quality within the window falls below a threshold of 15.

#### Differential expression analysis

Mapping was conducted independently for each species using RSEM (1.3.3; B. Li & Dewey, 2011) with the STAR (2.7.11a; Dobin et al., 2013) mapping to each respective reference genome (Supplementary material). The read counts were imported to R using tximport (3.19; Soneson C et al., 2015) within the R statistical computing environment (4.3.2; R Core Team, 2024) to avoid potential biases introduced by the differing lengths of isoforms, as this method employs estimated gene counts based on mapped transcript quantification in RSEM. The differential expression analysis was conducted using DESeq2 (1.12.3; Love et al., 2014) with a model based on the experimental design, larvae were compared to adults (male and female) using the pairwise comparison setting. To identify differentially expressed genes, a false discovery rate (FDR) of 0.05 was applied using the Benjamini-Hochberg method (Benjamini & Hochberg, 1995), and genes exhibiting a log2 fold change greater than 1 or less than −1 were considered to be significantly differentially expressed for each comparison. Each pairwise comparison yielded a list of genes more highly expressed in that condition, which were then used for GC content estimation (Table 1). Sample clustering was visualized with a principal component analysis (PCA) using DESeq2 on rlog-transformed data. This was done to assess the quality of the data, the overall effect of sex and stage on the dataset, and to rule out potential outlier samples.

**Table 1.** Summary of GC content intra-species comparisons across differentially expressed genes in larvae and adults.

| Order | Species | Comparison (number of DE genes) | Difference | Methylation | Mean GC content |
| --- | --- | --- | --- | --- | --- |
| Diptera | *Drosophila melanogaster* | Adults (3887) < Larvae (3796) | 4.88% | No | 48.49% |
| Diptera | *Bactrocera dorsalis* | Adults (3487) < Larvae (2078) | 1.63% | Yes | 40.69% |
| Diptera | *Belgica antartica* | Adults (1836) > Larvae (1439) | 1.81% | No | 38.87% |
| Hymenoptera | *Athalia rosae* | Adults (3472) < Larvae (5456) | 0.35% | Yes | 42.29% |
| Hymenoptera | *Lysiphlebus fabarum* | Adults (5201) < Larvae (4317) | 1.7% | No | 29.79% |
| Lepidoptera | *Manduca sexta* | Adults (3865) < Larvae (4286) | 8.96% | Yes | 35.65% |

#### GC content analysis

The fasta sequences of differentially expressed genes were retrieved using blastdbcmd of BLAST+ (2.14.1; Camacho et al., 2009) on the CDS version of reference genomes (Supplementary material). The GC content of the differentially expressed genes was calculated using seqinR (4.2-36; Charif & Lobry, 2007). The distribution and mode of the GC content were then measured and visualized using ggplot2 (3.5.1; Wickham, 2016), ggridges (0.5.6; Wilke C, 2024), and paletteer (1.6.0; Hvitfeldt E, 2021). Pairwise Wilcoxon rank sum tests (Mann & Whitney, 1947) were conducted to statistically compare the GC content distributions between all pairs of conditions. Subsequently, a comparison was conducted between the GC content distributions of the differentially expressed genes and those of all other genes. Any outliers identified in the differentially expressed genes were considered potential candidate genes for a GC shift between adults and larvae.

#### GC bias in orthologous genes

To ascertain whether the same genes were involved in the GC content bias during development, the orthologous genes present in six of the studied species were inferred using OMA (Orthologous MAtrix) (v2.5; Zahn-Zabal et al., 2020). This was done based on the protein sequences of the longest transcript of each gene, to avoid isoforms, and using *Manduca sexta* as the outgroup. The genomic sequences of the OMA genes were retrieved by blasting the protein id of OMA genes to the reference genome, and the GC content was subsequently measured using a customed script. The distributions of GC content in OMA genes and differentially expressed genes were compared using the Wilcoxon rank sum test. Outliers in the group of differentially expressed genes were extracted using the interquartile range (IQR) method in R, and those present in multiple comparison per species represent candidate genes for GC shift, with the OMA-only genes used as a reference. Candidate genes shared between 2 or more species are presented in the Supplementary Table S1.

#### Gene ontology enrichment analysis

Stage-biased gene sets (larval-biased and adult-biased) in *Drosophila melanogaster* were analyzed for functional enrichment of biological process categories using enrichGO function from clusterProfiler (4.16.0; Yu et al., 2024) with the org.Dm.eg.db annotation package (*Drosophila melanogaster*, FlyBase gene identifiers). *Drosophila melanogaster* being a biological model, information allowing us to do the enrichment analysis were available, which is not the case for the five other species. Significance was defined as adjusted p-value < 0.05 with the Benjamini-Hochberg method. The top 15 enriched GO terms per stage were displayed using dot plots, with point size proportional to the number of associated genes and color reflecting enrichment significance. To explore functional redundancy among GO terms, pairwise semantic similarity was conducted using pairwise_termsim function with default Jaccard correlation coefficient, and results were visualized with the ssplot function from enrichplot (1.28.4; Yu et al., 2025). Node labels were shown at the category level.

### Whole-genome GC content through insect orders

To investigate the influence of taxonomic order and parasitic behavior on the mean GC content of coding sequences from 317 insect genomes, we built two binomial generalized linear models (GLMs) with a logit link function, as GC content is expressed as a proportion. The first model was structured as: %GC ∼ Order + Behavior, and the second as: %GC ∼ Order × Behavior.

The first model, which excluded interaction terms, was applied to all 317 species, encompassing 22 Coleoptera, 102 Diptera, 122 Hymenoptera, and 71 Lepidoptera. Of these, 222 species were non-parasitic and 95 were parasitic. This analysis was conducted using the *stats* (v4.2.3; R Core Team, 2024) and *sjPlot* (v2.8.16; Lüdecke, 2024) R packages.

The second model, which included an interaction term, was restricted to Diptera and Hymenoptera. Only these orders had sufficient representation of both parasitic and non-parasitic species to investigate the interaction between order and parasitic behavior. Odds ratios (representing the odds for species to have a higher GC content compared to a reference) derived from both models are presented in the results section.

To further investigate biases across a wide range of genes, codon usage was assessed using relative synonymous codon usage (RSCU). The RSCU values were calculated from the complete set of annotated coding sequences (CDS) for the six species used for the RNA-seq analysis, using the *seqinR* package (v4.2-36; Charif & Lobry, 2007) in R. These values were visualized with *ggplot2* (v3.5.1; Wickham, 2016) and further refined using *Inkscape* (v1.3.2; Inkscape Project, 2023).

## Results

### GC content across insect orders

To examine the influence of behavior and taxonomic order on the GC content at the genome-level, we built a GLM using CDS from 317 species with: 22 Coleoptera, 102 Diptera, 122 Hymenoptera, and 71 Lepidoptera. Of these, 222 species were non-parasitic and 95 were parasitic. A statistically significant influence of insect order on GC content was found by the GLM evaluating the GC content across taxonomic groupings and lifestyles (p < 0.001; Fig. 1A; S2, S3), demonstrating significant variation among various orders. With odds ratios (OR) of 1.29 and 1.33, respectively, species from the orders Diptera and Hymenoptera had the greatest mean GC content when compared to the reference order Coleoptera (p < 0.001; Fig. 1A). Additionally, with an odds ratio of 0.76 (p < 0.001; Fig. 1A), parasitic species tended to have lower GC content than non-parasitic species.

**Figure 1.**
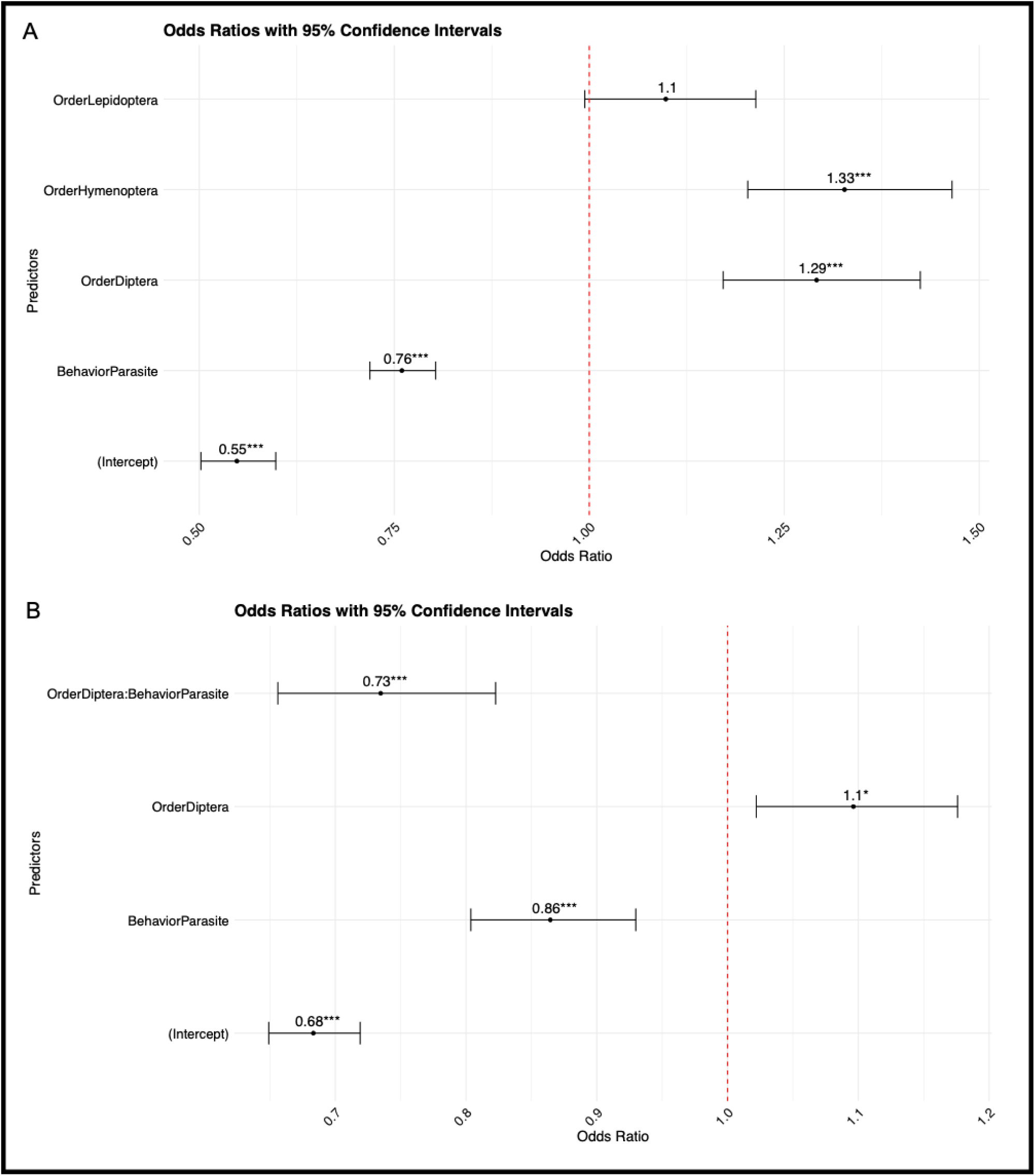
Odds ratio of linear model for GC content analysis across insect orders and parasitic behavior. This figure shows the odds ratios and 95% confidence intervals for the effect of taxonomic order and parasitic behavior on GC content from two GLMs. **A**. In the first model, higher GC content in Diptera and Hymenoptera compared to Coleoptera (p < 0.001) and lower GC content in parasitic species (p < 0.001) were observed. **B**. The model with interaction shows that parasitic Diptera and Hymenoptera have lower GC content (OR = 0.73 and OR = 0.86 p < 0.001). Significance levels are indicated by asterisks (*p < 0.05, ***p < 0.001) and the odds ration threshold by a red dashed line.

The results of the second model (included an order-lifestyle interaction) also identified a significant decrease in GC content in parasitic species. With an odds ratio of 1.10 (p = 0.010; Fig. 1B, S3), Diptera species had a higher mean GC odds ratio than the reference order Hymenoptera, even though Hymenoptera showed higher GC content in the first model. Furthermore, with an odds ratio of 0.86 (p < 0.001; Fig. 1B), parasitic species had a reduction of GC content compared to non-parasitic species. Essentially, the interaction term showed that the high GC content seen in non-parasitic Diptera was offset by a much lower GC content in parasitic Diptera, with an odds ratio of 0.73 (p < 0.001; Fig. 1B). Divergent trends in GC content are observed within the same taxonomic order, which can impact the overall patterns seen at the order level. In summary, the two statistical models identified the substantial influence of both taxonomic order and parasitic lifestyle on GC content in insects. Further investigation is necessary to elucidate the mechanisms underlying these variations.

### GC shifts during development

To examine life-stage associated GC content shifts, we utilized available transcriptomic data from adult and larval stages in six species: two Hymenoptera (*Athalia rosae* and *Lysiphlebus fabarum*), three Diptera (*Drosophila melanogaster, Bactrocera dorsalis, Belgica antartica*), and a Lepidoptera (*Manduca sexta*). In the six tested species, mean GC content across all coding genes varied between species, with a wider range (29.79% to 42.29%) observed in Hymenoptera compared to Diptera (38.87% to 48.49%), Table 1).

Among differentially expressed genes, the GC content comparisons between developmental stages revealed that, in five out of the six species, the differentially expressed genes exhibited a higher GC content in larvae (Figure 2). A Wilcoxon rank sum test indicated that the GC distribution in larvae and adults is significatively different across all species (Figure 2). The only exception was in Diptera, where GC content in differentially expressed genes was higher in larvae in *Bactrocera dorsalis* and *Drosophila melanogaster* (Fig. 2A, 2B), but not in *Belgica antartica* (Fig. 2C). Higher GC content in larvae was observed in both hymenopteran species (Fig. 2D, 2E) and in *Manduca sexta* (Fig. 2F), revealing a pattern of GC bias between developmental stages. Looking at the median GC content in both developmental stages, the difference between larvae and adults ranges from 0.35% to 8.96% with the greatest differences of 4.88% in *Drosophila melanogaster* and 8.96% in *Manduca sexta* (Tab. 1).

**Figure 2.**
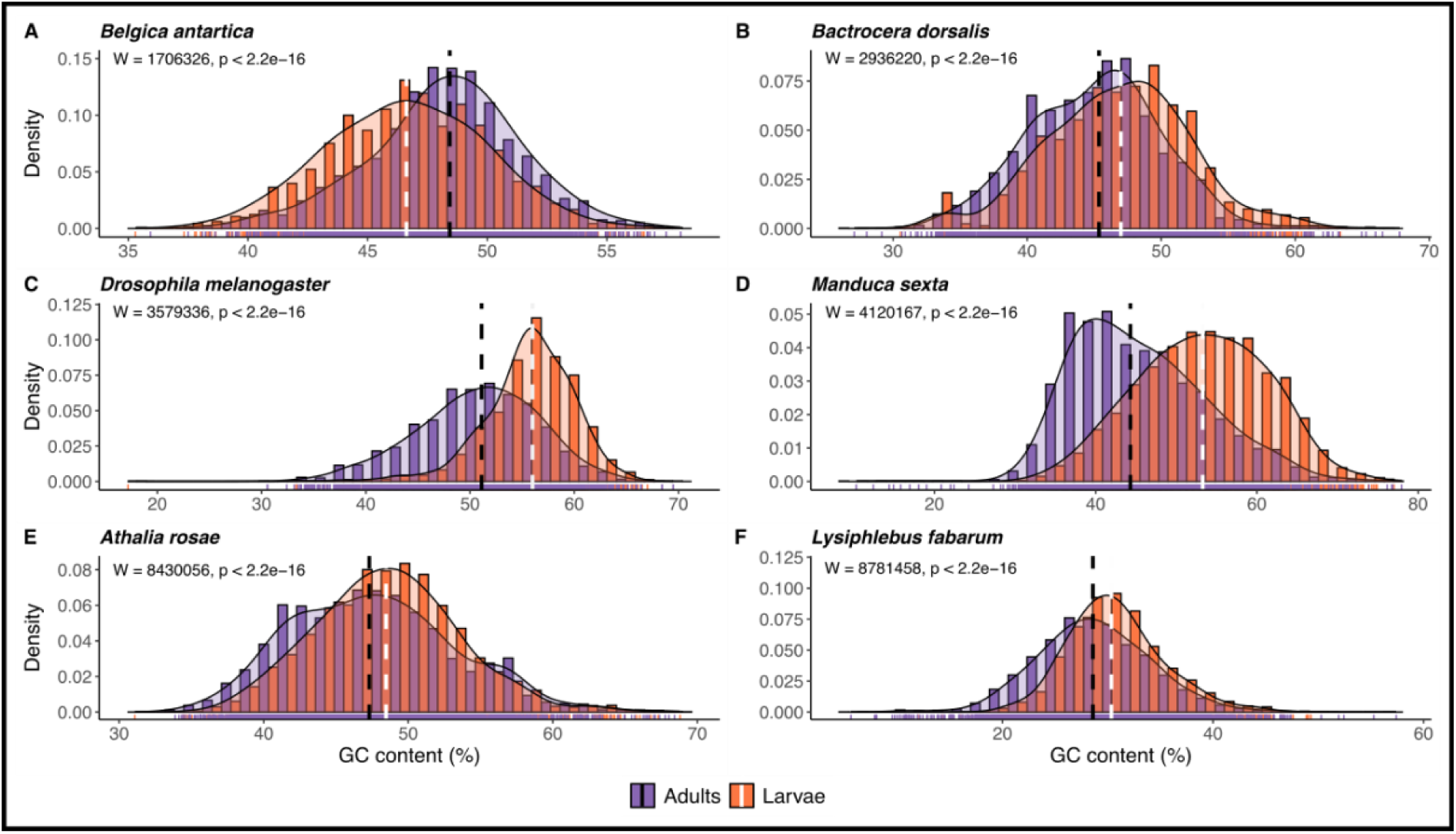
Distribution of GC content in expressed genes of insect species across different life stages. This figure shows the GC content distribution in larvae and in adults in Diptera (**A-C**), Lepidoptera (**D**) and Hymenoptera (**E-F**) orders. Statistical tests used was Wilcoxon rank sum test.

#### Adult-biased genes are reproduction related

To characterize the functional roles of stage-biased genes, a GO enrichment analysis was performed on larvae- and adult-biased differentially expressed genes in *Drosophila melanogaster*.

Adult-biased genes were enriched in cilium movement, microtubule-based movement and microtubule cytoskeleton organization, but also in reproduction related processes as sperm motility and eggshell formation (Fig. 3B). A semantic similarity plot revealed clustering of related genes category such as axonemal formation complex, egg formation, male gamete formation, and motility and fecundation related processes (Supplementary material). In contrast, larvae-biased genes are significantly enriched in cytoplasmic translation, oxoacid metabolic process, carboxylic acid metabolic process and organic acid metabolic process, as well as cuticle development and catabolic processes (Fig. 3A). A semantic similarity plot revealed clustering of related gene ontology categories, highlighting that larval-enriched terms forms cohesive groups related to development and biosynthetic pathways with an enrichment in catabolic, metabolic and translation processes (Supplementary material).

**Figure 3.**
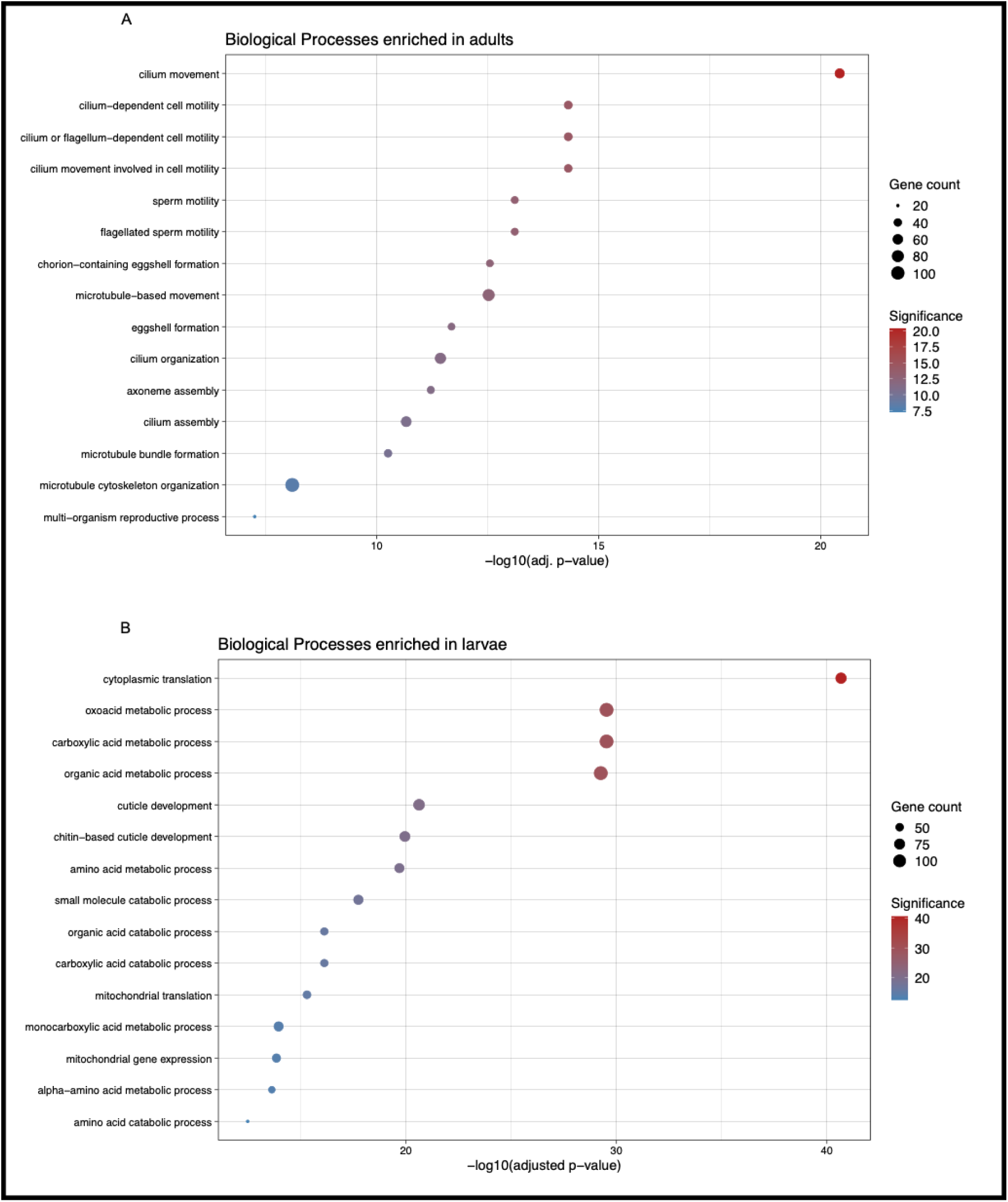
Gene ontology enrichment of biological processes in stage-biased genes in *Drosophila melanogaster.* Dot plot of adult-biased DEGs showing significant enrichment in cilium movement, sperm motility and egg formation (A). Dot plot of larvae-biased DEGs showing enrichment in cellular chitin-based cuticle development, translation, catabolic, and biosynthetic processes (B).

In summary, adult-biased genes are generally enriched in reproductive processes and larval-biased genes in development and energy production through biosynthetic processes.

#### Orthologous gene comparison

Given that we have demonstrated GC content shifts across life stages, our next objective was to ascertain whether this observed GC content bias is attributable to the presence of shared differentially expressed genes among the species under investigation.

Using the predicted coding sequences from the six species’ genome annotations, we compared GC content distributions between differentially expressed and non-differentially expressed genes within those that were identified as one-to-one orthologous genes. In the species of Diptera, GC content in differentially expressed and non-differentially expressed orthologous genes was significantly higher in *Bactrocera dorsalis* (MW: W = 1189081, p < 0.001; Fig. 4A), *Belgica antartica* (MW: W = 1327742, p < 0.001; Fig. 4B) and *Drosophila melanogaster* (MW: W = 1771392, p-value < 0.001; Fig. 4C). In Hymenoptera, notable discrepancies in GC content were detected in *Athalia rosae* (MW: W = 2466618, p < 0.001; Fig. 4E) with higher GC content observed in differentially expressed genes, but not in *Lysiphlebus fabarum* (MW: W = 3768207, p = 0.7153; Fig. 4F). Finally, in representation of Lepidoptera, GC content was significantly lower in *Manduca sexta* orthologous genes compared to non-differentially expressed orthologs (MW: W = 1747950, p < 0.001; Fig. 4D). A pattern seems to emerge from differentially expressed and non-differentially expressed orthologous genes GC content comparisons across species, with higher GC content in differentially expressed genes. Moreover, outliers exist in differentially expressed genes and can be investigated to identify potential adaptations.

**Figure 4.**
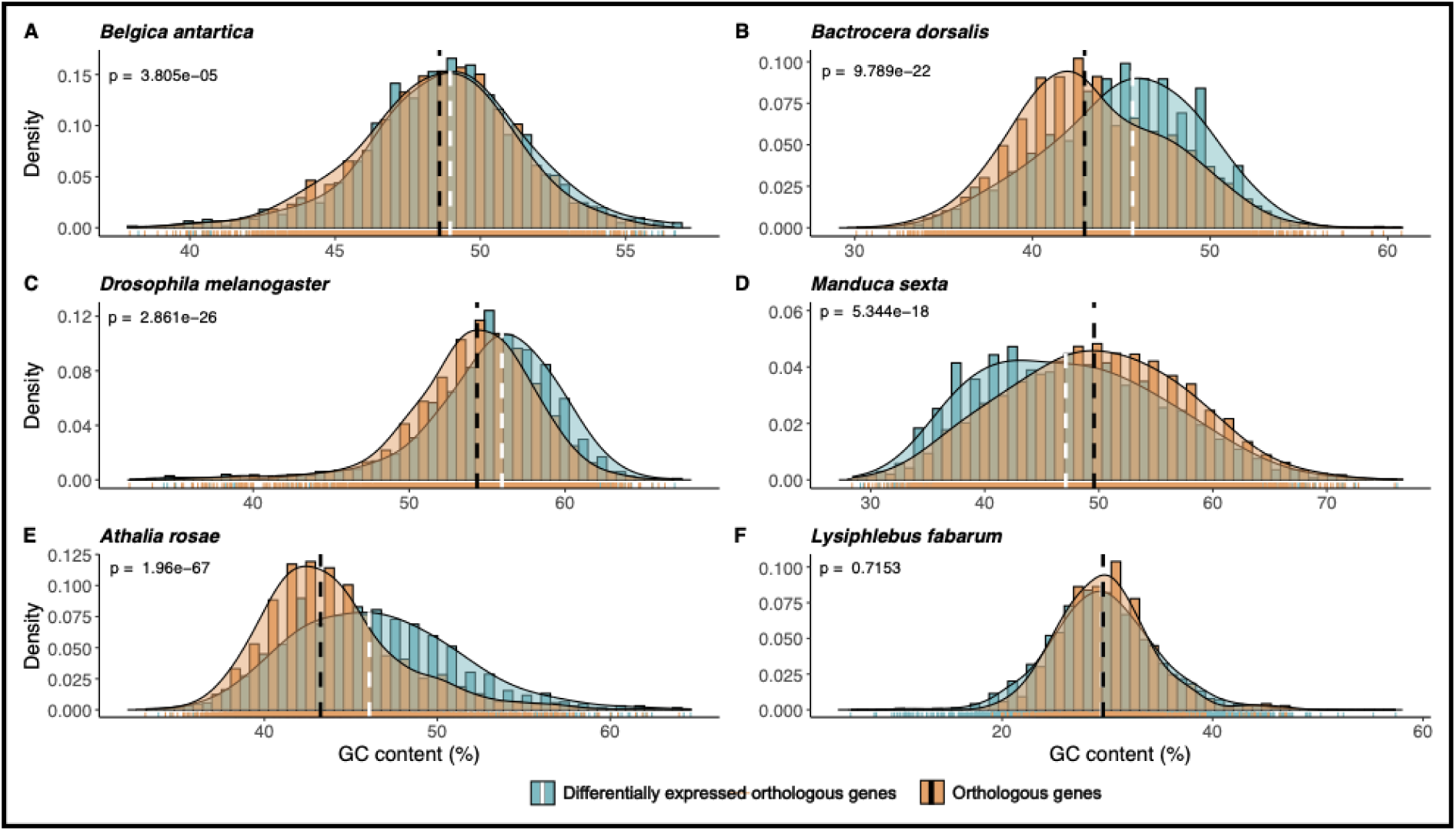
Distribution of GC content among orthologous genes and differentially expressed genes that are shared across species. The GC content of differentially expressed genes is observed to be higher to that of orthologous genes in Diptera (A, B, C) and markedly higher in *Athalia rosae* (E) but comparable in *Lysiphelbus fabarum* (F). In Lepidoptera species, the distribution of GC for differentially expressed genes (D) is observed to be lower.

Following this overall comparative analysis, outliers in differentially expressed orthologous genes were identified by blast.

In total, we identified 18 candidate genes that were shared between 2, 3 or 4 species (see Supplementary Table, S1). An example is the “glycin-rich cell wall structural protein 1.8 gene”, which is shared between *Athalia rosae*, *Belgica antartica, Drosophila melanogaster* and *Lysiphlebus fabarum*, with GC contents of 61.03%, 55.37%, 67.02% and 41.29%, respectively. This gene is expressed in both larvae and adults, depending on the species, and contributes to the molecular structure of the cuticule. Another common candidate is the probable H/ACA ribonucleoprotein complex subunit 1 gene shared by *Athalia rosae*, *Belgica antartica* and *Lysiphlebus fabarum* with 52.57%, 55.91% and 44.63% of GC, respectively. It is expressed preferentially in larvae *Belgica antartica* and *Lysiphlebus fabarum* and contributes to ribosome biogenesis and telomere maintenance. Other genes were involved in cell structure and energy production (S1).

## Discussion

Our analysis of GC content variation across insect genomes showed that mean GC content at the whole genome-level varied in relation to multiple factors, including taxonomic order and parasitic lifestyles. Moreover, we showed that GC content differed between developmental stages within species, even though the genome remains the same. Both suggest multiple evolutionary pressures on GC content, but at different levels, possibly driven by multiple mechanisms.

### Gene expression and translation efficiency as drivers of GC variation

Our analysis revealed that, in five of the six species tested, larvae exhibited a higher GC content in their differentially expressed genes compared to adults. A slight trend was observed in the Hymenoptera and Lepidoptera orders. Conversely, in the Diptera order, two of the three tested species exhibited higher GC content in adult-biased genes compared to larvae. While further validation is required using higher-quality samples and a more extensive range of species within each order, these findings imply a potential shared transcriptomic bias in GC-content across insect species. To better understand the reasons why GC content can vary depending on the stage and species, a better understanding of gene expression regulation is needed. We suggest that stability of the mRNA, affecting translational efficiency and the codon usage, linked to deficiency of methylation in insect, are a potential answer to that variation, and should be better studied.

Multiple factors can influence variation in GC content across genomes and developmentally stage-biased genes. Among these are the mechanisms that regulate gene expression, most notably methylation. Methylation has been identified as a critical process that contributes to shaping the GC landscape of the genome. When methylation is absent or reduced, there may be less selective pressure for high GC content in the upstream gene areas (Bewick et al., 2016; Mugal et al., 2015). For example, while mammals exhibit elevated GC content in the 5’- and 3’-UTRs compared to CDS, studies in *Drosophila melanogaster* and *Caenorhabditis elegans* reveal the opposite trend, with coding regions exhibiting higher GC content than in UTRs (Zhang et al., 2004). Lower GC content in the 5’-UTR in these species could be a consequence of their loss of methylation, while the higher GC content in CDS might result from the necessity to maintain mRNA stability. Similarly, methylation may also provide an explanation for the shifts in GC content that we have observed across different developmental stages. An explanation could be that genes expressed in juvenile tissues may exhibit elevated GC content due to methylation. For example, in mammals, methylation of CpG sites functions as a regulatory mechanism for gene expression, serving as a switch to fine-tune gene activity across tissues and developmental stages and leading to high GC content (Hapgood et al., 2001). However, several insect orders, including Diptera, some Hymenoptera, and Lepidoptera, exhibit absent or reduced methylation levels at early developmental stages (Bewick et al., 2016). Lack of methylation may reduce the need for GC-rich areas, which would lessen the selective pressure that would otherwise be applied to maintain these energetically expensive nucleotides. Another potential factor to consider is the lower recombination rate in some genomic regions, which disrupts the equilibrium between AT and GC-biased mutations, favoring the accumulation of AT-rich genes across the genome (Kotari et al., 2024).

Additionally, it has been observed that genes with a high GC content generally show less stable mRNA around the translation-initiation site. This phenomenon has been shown to enhance translation efficiency by facilitating ribosome accessibility (Gu et al., 2010). Synonymous mutations, which affect translation efficiency by influencing mRNA degradation, could also play a role. Guanine and cytosine, with their three hydrogen bonds, have been shown to confer greater stability to mRNA compared to adenine and uracil. This increased stability may help buffer mRNA against environmental variations such as thermal fluctuations (Liao et al., 2021). Although mRNA stability does not appear to directly influence gene expression in *Drosophila melanogaster* (Eck & Stephan, 2008), it could still be associated with high GC content in stage-biased genes. This relationship may reflect a balance between global mRNA stability and the need for rapid translation. In our study, depending on the species, the median GC shift ranged from 0.35% to 8.96% between stage-biased genes. Such shifts have the potential to influence developmental stages, as discussed previously, the elevated need for translation efficiency in larvae may drive the evolution of genes with higher GC content in these stages, as they need to produce proteins for their rapid growth at a high rate in holometabolous to complete metamorphosis. The holometabolous strategy was shown to remove developmental constraints like the fast growth – differentiation trade-off, and reduce mortality (Manthey et al., 2024).These benefits are particularly important during larval development, where rapid growth and elevated energy demands are paramount.

While the stability of mRNA is critical to the maintenance of high GC content in coding sequences, codon usage also significantly influences gene regulation and translation efficiency. Optimal codon usage has been demonstrated to enhance translation elongation speed in *Drosophila melanogaster*, exhibiting a slight bias toward GC-rich codons (GC3 = 64.32%) relative to AT-rich codons (Powell & Moriyama, 1997; Zhao et al., 2017). This predilection for GC-rich codons is believed to enhance translational efficiency by facilitating accelerated elongation rates, a process that is particularly crucial for rapidly transcribed genes, as it might be necessary during larval development. And even more so in parasites. It is noteworthy that codon usage bias is not exclusively associated with high GC content. For example, *Lysiphlebus fabarum*, a species with a GC-poor genome, also exhibits codon usage bias (Dennis et al., 2020). Other codon bias studies revealed that insects exhibited base composition preferences. In dipterans, codons ending in G or C were observed to be preferred, whereas in hymenopterans, codons ending in A and T were found to be preferred (Behura & Severson, 2013). This finding indicates that codon usage bias, rather than the overall GC content, plays a substantial role in the optimization of translation in organisms with diverse genomic GC content. Indeed, codon usage can be a pivotal element in facilitating efficient protein synthesis, irrespective of the genome-wide GC content.

We also observed that adult-biased genes were enriched in biological processes related to reproduction such as gamete production, sperm motility and fecundation. In contrast, larval-biased genes were enriched in developmental and energy production related functions. Even if there is a higher energy requirement in females for processes such as egg production and offspring nourishment (Ingleby & Morrow, 2017), there is more investment in the production of energy in larvae to sustain the development. GC-rich regions have been demonstrated to support increased gene expression and stability in order to meet these demands, in females and larvae.

Altogether, these observations underscore the correlation between GC content variation and developmental and sex-specific biological imperatives. High levels of GC content in specific stages or sexes may be indicative of a requirement for expeditious and effective gene expression to support critical physiological processes, including growth, development, and reproduction.

### Parasitism as a driver for low GC in insects at genome-level

Across 317 species of insects, we examined GC content as a function of taxonomic order and behavior (model: GC ∼ Order + Behavior). This highlights the significant roles of both taxonomic group and lifestyle in driving GC content evolution in insects, specifically in reducing GC content in parasitic species. This finding is further substantiated by the prevalence of AT-rich genomes in ciliates, fungi, apicomplexans, and amoeba, with some of these organisms displaying parasitic behaviors (Videvall, 2018). Among 81 species of Platyhelminthes and Nematoda, 72% display a mean genomic GC content below 40% (International Helminth Genomes Consortium, 2019). The low GC content observed in parasitic species can be attributed to reduced recombination rates and AT-biased mutations, or even to parasite-specific patterns of expression, as has been documented in *Trypanosoma* (Martinez et al., 2014). AT-biased mutations have been observed in intracellular eukaryotic microbes based on bacterial analogs (Videvall, 2018). However, further research is needed to elucidate the precise mechanisms underlying this nucleotide composition bias.

Parasites frequently encounter nutrient limitations, necessitating the implementation of strategies to minimize metabolic expenditure. One such strategy may involve genomic streamlining to conserve energy, as reflected in the elimination of GC-rich regions in parasitic platyhelminths (Chen et al., 2013). While GC-rich codons are associated with lower energy costs for amino acid production (Seligmann, 2003), AT-rich codons may reduce the energetic burden of nucleotide synthesis (Bohlin et al., 2013; Kim et al., 2016). This tradeoff enables parasites to allocate their resources toward amino acid production rather than nucleotide synthesis, thereby facilitating their survival under nutrient-constrained conditions. Furthermore, some parasitoid insects *Lysiphlebus fabarum*, have undergone a loss of the capacity to synthesize specific amino acids. Consequently, these insects are reliant on their hosts’ amino acid pools and protein degradation pathways to satisfy their nutritional needs (Ye et al., 2022). This metabolic dependency may further conserve energy for essential biological functions, such as growth and development.

A further potential contributor to the reduced GC content observed in parasite genomes is the prevalence of transposable elements. These sequences are predominantly AT-rich, although their nucleotide composition can vary (Boissinot, 2022; Ruggiero & Boissinot, 2020). The increased activity of transposable elements in parasite genomes could have the potential to amplify AT-rich regions, thereby further driving the observed reduction in GC content (Boissinot, 2022). The investigation of the proportion of repetitive sequences in insect genomes could elucidate their contribution to GC content variation across different lifestyles.

The interplay of these factors—AT-biased mutations, reduced recombination, genomic streamlining, reliance on host-derived nutrients, and transposable element prevalence—has been hypothesized to underlie the low GC content observed in certain parasitoid species, such as *Anastatus japonicus* and *Anastatus fulloi* (Ye et al., 2022). Future studies focusing on the interactions among these processes will provide deeper insights into the evolutionary and ecological drivers of GC content variation in parasitic insects.

### Mosaic genome, synonymous mutations and selection

This study, together with existing literature, highlights the variability of GC content across genomes and among species, emphasizing its evolutionary adaptation to diverse environments over generations. As previously discussed, the observed differences in GC content are indicative of distinct mechanisms and selective pressures acting on genes (Pozzoli et al., 2008). However, it should be noted that GC content variability exerts a significant influence not only on organismal biology but also on analytical approaches. For instance, our findings exemplify how multiple factors at various levels contribute to GC variation, complicating its interpretation and introducing biases. Romiguier and Roux (2017) detailed these biases, which range from challenges in constructing phylogenetic trees to estimating codon usage, complicating studies of evolutionary processes (Romiguier & Roux, 2017).

The detection of selection is a good example of this phenomenon. Current methods frequently are based on the assumption that synonymous mutations are neutral. However, this assumption has been refuted; researches have demonstrated the influence of selection on synonymous mutations, under negative selection (Gudkov et al., 2024) and positive selection (Poon et al., 2021), showing the necessity to reevaluate this framework. Next-generation sequencing development have rendered feasible comprehensive genomic and transcriptomic analyses, thereby enabling more precise investigations of biological phenomena associated with GC content variation. These resources provide a way to clarify the nuances of adaptation and selection as they appear in GC content.

### Meta-analyses rely on data quality

The findings of this study are contingent upon the quality of the data that has been re-used for this study. The results of this are dependent on the integrity of the data and reliability of the sampling methods of original researchers. The table of reference genomes indicates the developmental stage used to construct the reference genome, which can vary from larval to adult and between male and female. For instance, the *Bactrocera dorsalis* reference is an adult female, which may impede the aligner’s capacity to map male-specific reads to this genome, this is a sex-dependent problem if the reference genome is not supplemented with the other sex-specific chromosomes. It is possible that the data may be influenced by factors such as age variation within a stage, as well as by external variables such as stress, diet and sampling times. However, these variables are not always mentioned in the papers used for this meta-analysis. It is also important to note, that a reference genome may prove a suitable match for some samples but not for others (Monnahan et al., 2020), as the data quality and mutational differences between the sequenced populations might play a role in reads mapping. These differences can also be associated to the misannotation of reference genomes. Until now, new assembled genomes were annotated using one of the available softwares and protein were not experimentally characterized, leading to annotation errors in molecular function of enzymes (Schnoes et al., 2009). More alarming, misannotation of one conserved portion of 23S rRNA from NCBI genomes generated 90% of false positive matches protein in metatranscriptomics studies (Tripp et al., 2011). However, this situation is evolving with better annotation method using Hidden Markov Model and deep learning, leading to accurate annotation in mammals (Gabriel et al., 2024). This model needs to be trained on non-vertebrate genomes to allow *de novo* assembly and annotation in the future.

A further crucial element in meta-analysis is the size of the sample employed in the studies that we have re-analyzed. The sample size serves as a safeguard to ensure the reproducibility of the study, the generalization of the results to a species, and to establish an ethical aspect by striking a balance between precision and the number of animals used (Lakens, 2022). In this study, six species were utilized from a multitude of sources. While some of these sources provide information regarding the sample size, others do not, resulting in species with unknown sample size, this is the case for data coming from project without an associated paper. This could be problematic if the sample size were insufficient, allowing biological variability to predominate in the observed effects as the model fitting compares variation within conditions against the variation between conditions. However, the studies included in this meta-analysis have already undergone a rigorous peer-review process, ensuring the validity of the findings.

## Conclusion

In this study, we conducted a comprehensive analysis of transcriptomic sequences from larvae and adults across seven insect species and identify GC shifts associated with developmental stages. The results of this study suggest a potential shared selective pressure among insect species from three orders. We postulate several explanations for the observed patterns of GC bias, including the roles of gene regulation mechanisms and translational efficiency, as well as more broad-scale genomic evolution. However, further investigation is required to confirm these hypotheses.

From a technical perspective, our approach demonstrated the capacity to discern shared patterns of genomic evolution across species, separate from conventional selection tests. GC bias, often associated with codon usage bias, is influenced by synonymous mutations, which are typically considered to be neutral in selection tests. However, the evidence that we present here indicates that synonymous mutations can have important evolutionary consequences, possibly influencing gene regulation, mRNA stability, and translation efficiency. These findings challenge the assumptions underlying current selection tests and highlight the need for complementary approaches. These do not invalidate current methods but rather suggest that there is a layer of evolutionary change that they do not consider. Advancements in next-generation sequencing and the expansion of genomic databases present a promising avenue for a more comprehensive understanding of molecular evolution and adaptation.

## Supporting information

Data accession tables

## Acknowledgments

We thank Stefanie Hartmann for conducting the OMA analysis on the HPC Cluster of the University of Potsdam.

