## Supplemental figures S2-S4 for "Identifying transcriptomic bias across developmental shifts in insects"

### Supplementary files

**Table S1. Candidate genes shared across tested species summary.**
Table_S1_candidate_genes.xlsx

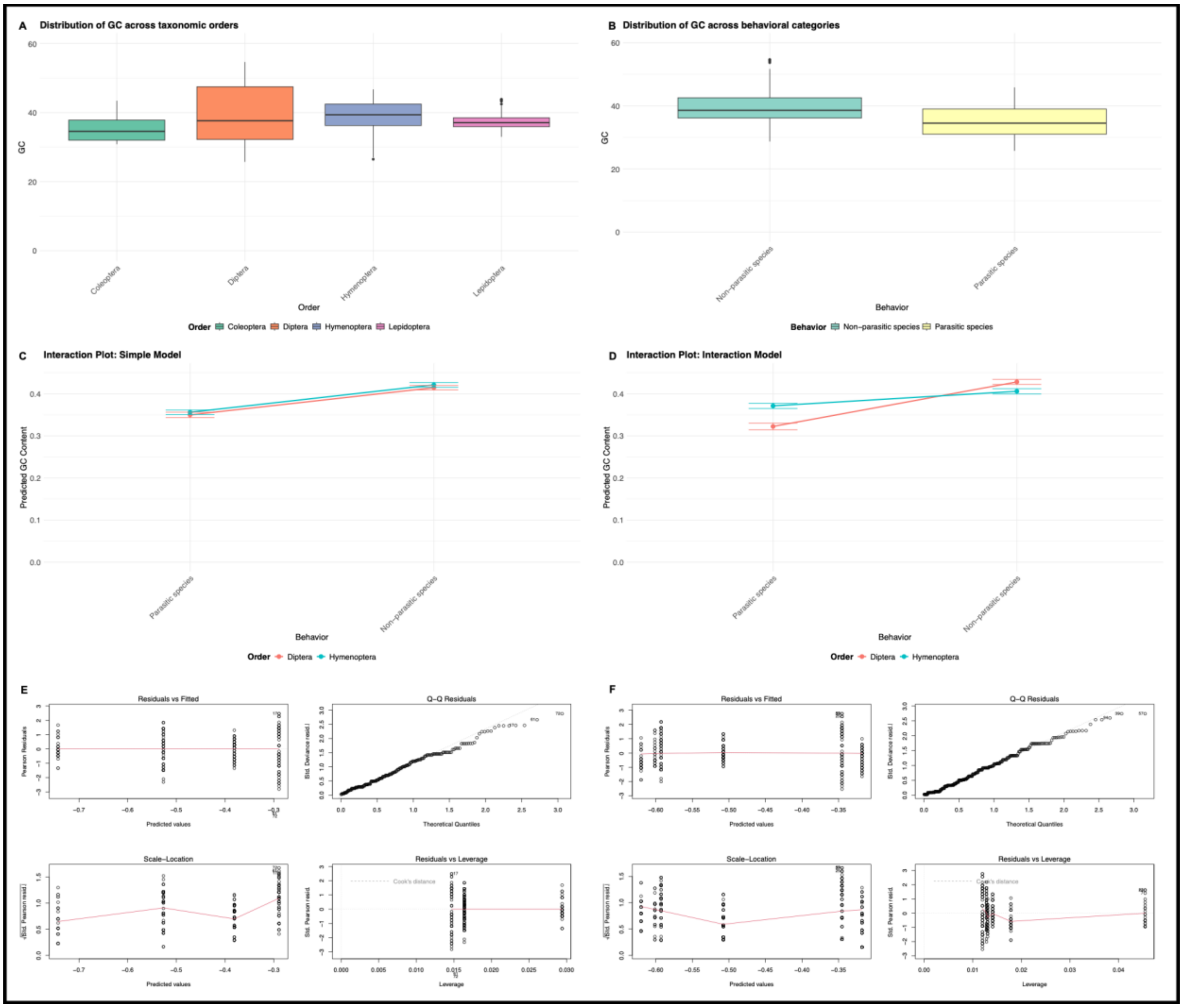


**Figure S2. Diagnostic graphs of the linear models. A.** Mean GC content at genome-level distribution across insect orders. **B.** Mean GC content at genome-level distribution across parasitic and non-parasitic insects. **C.** Interaction plot illustrating the effect of behavior and order in the simple model. **D.** Interaction plot illustrating the effect of behavior and order in the interaction model. **E.** Diagnostic plots for the simple model, showing residuals and model fit. **F.** Diagnostic plots for the interaction model, highlighting residuals and fit quality.

A

| logit(P(GC content))=β0+β1(Order)+β2(Behavior) | | | |
|---|---|---|---|
| *Predictors* | *Odds Ratios* | *Conf. Int (95%)* | *P-Value* |
| Coleoptera | *Reference* | | |
| Diptera | 1.29 *** | 1.17 – 1.42 | **<0.001** |
| Hymenoptera | 1.33 *** | 1.20 – 1.47 | **<0.001** |
| Lepidoptera | 1.10 | 0.99 – 1.21 | 0.066 |
| Non_parasite | *Reference* | | |
| Parasite | 0.76 *** | 0.72 – 0.80 | **<0.001** |
| Observations | 317 | | |

** p<0.05 ** p<0.01 *** p<0.001*

B

| logit(P(GC content))=β0+β1(Order)+β2(Behavior)+β3(Order×Behavior) | | | |
|---|---|---|---|
| *Predictors* | *Odds Ratios* | *Conf. Int (95%)* | *P-Value* |
| Hymenoptera | *Reference* | | |
| Diptera | 1.10 * | 1.02 – 1.18 | **0.010** |
| Non_parasite | *Reference* | | |
| OrderDiptera:BehaviorParasite | 0.73 *** | 0.66 – 0.82 | **<0.001** |
| Parasite | 0.86 *** | 0.80 – 0.93 | **<0.001** |
| Observations | 224 | | |

** p<0.05 ** p<0.01 *** p<0.001*

**Figure S3. Summary of the linear models explaining mean GC content in the genomewith the behavior and Order of insects. A.** Summary of the linear model without interactions. **B.** Summary of the linear model with interactions.

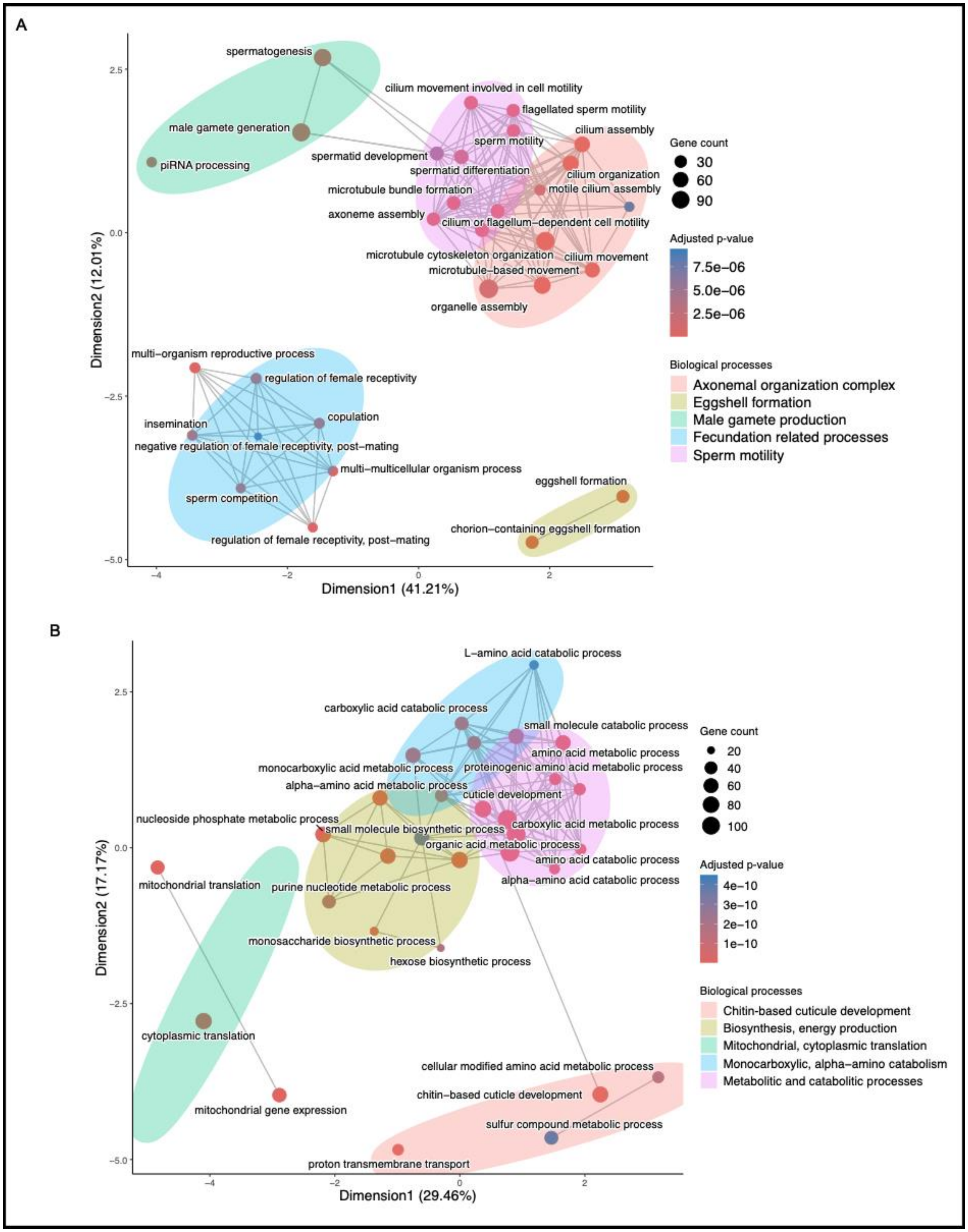


**Figure S4. Gene ontology analysis in *Drosophila melanogaster* for adults and larvae.**
**A.** Semantic plot of the enriched terms in adults. **B.** Semantic plot of the enriched terms in larvae.

Supplementary_material_Data_accession.xlsx
